# Effector-triggered stomatal immunity prevents leaf invasion by bacterial pathogens

**DOI:** 10.64898/2026.03.09.710543

**Authors:** Tamar V. Av-Shalom, Yan Lai, Racquel A. Singh, Dora Yuping Wang, Reid Gohmann, David Mackey, Darrell Desveaux, David S. Guttman

## Abstract

The phytopathogen *Pseudomonas syringae* must enter the leaf interior through natural openings such as stomata to cause disease. Plants restrict pathogen entry into the leaf upon non-self recognition-driven closure of their stomata. *P. syringae* counters this stomatal immunity by deploying a range of toxins and effectors. Here, we show that stomatal immunity in *Arabidopsis thaliana* is reinforced upon recognition of the ubiquitous *P. syringae* type III effector AvrE1 by the resistance protein CAR1. *CAR1* expression is enriched in guard cells, and its activation prolongs the effective duration of stomatal immunity. We also show that CAR1 prevents the invasion of leaf tissue by natural isolates of *P. syringae*, demonstrating that CAR1-mediated stomatal immunity against the conserved effector AvrE1 provides a broadly significant barrier to host accessibility by *P. syringae*.

## Introduction

The plant immune system can be divided into two levels of microbial recognition. Pattern recognition receptors (PRRs) induce a basal level of immunity, referred to as Pattern-Triggered Immunity (PTI), upon recognition of microbe-associated molecular patterns, such as bacterial flagella ^1,2^. However, many phytopathogens have evolved arsenals of effectors that suppress PTI ^3^. In turn, plants trigger an amplified immune response, termed effector-triggered immunity (ETI), upon recognition of pathogen effectors ^4,5^. Typically, ETI is mediated by intracellular nucleotide-binding leucine-rich repeat immune receptors (NLRs) that are triggered when they directly bind effectors or indirectly detect effector activity within the plant cell ^5,6^. Although the two branches rely on distinct receptors for microbial perception, they are tightly interconnected with shared signaling components and mutual reinforcement of downstream responses ^4,7^

As plants lack migrating immune cells, it was historically speculated that all plant tissues are equally immunocompetent. However, recent studies utilizing single-cell sequencing have shown vastly different, tissue-specific patterns of immune activation against pathogen attack ^8,9^. Transcriptomic analysis of PTI responses in three different root cell types: epidermis, cortex, and pericycle cells, has shown distinct expression of immunity-related genes ^10^. Cell-specific responses to bacterial flagellin alter the ability of bacteria to colonize *A. thaliana* roots ^11^. While few examples of tissue-specific NLR-mediated responses have been identified to date, recent studies have revealed root and hydathode-specific immune responses. A cluster of tomato resistance genes against cyst and root-knot nematodes was recently found to be nearly exclusively expressed in the root tissue ^12^. Another study showed that the *A. thaliana* NLR, SUT1, was required for early hydathode immunity against the bacterial pathogen *Xanthomonas campestris* pv. *campestris*, but ineffective if the bacteria bypass hydathode entry ^13^. Finally, a complex between the helper NLR ADR1 and the lipase-like proteins EDS1 and PAD4 has been shown to mediate stomatal closure in PTI, although the expression of these proteins is not specific to the guard cells ^14^. Although these studies indicate tissue-specificity, an effector-triggered and tissue-specific immune response has yet to be identified.

Stomatal apertures are dynamically controlled during different phases of plant interactions with foliar bacterial phytopathogens, such as *P. syringae*. Initiation of infection requires invasion of the leaf apoplastic space through wounds or natural openings on the leaf surface, such as stomata ^15^. Stomatal immunity is an important early output of PTI that occurs when plants encounter microbes and close their stomata to prevent invasion^16^. Phytopathogens, such as *P. syringae*, have evolved toxins and type III secreted effector proteins that promote virulence by suppressing PTI, including stomatal immunity^3,17–19^. These include the phytotoxin, coronatine, and effectors, including AvrB1, HopF2 and HopX1, that specifically counter stomatal immunity ^16,20–23^. Once stomatal immunity has been breached, stomatal closure now benefits the pathogen rather than the plant host, since closed stomata reduce transpiration, and the resulting increase in apoplast hydration favors bacterial proliferation. *P. syringae* deploys functionally redundant effectors, HopM1 and AvrE1, each of which modulate ABA synthesis and/or trafficking to promote stomatal closure ^24–26^. Overall, the control of stomatal aperture is a crucial aspect of plant-pathogen interactions and a fundamental selective pressure driving the host-pathogen evolutionary arms race.

In this study, we show that ETI plays an important role in stomatal immunity. We find that the *A. thaliana* NLR, CEL-ACTIVATED RESISTANCE 1 (CAR1), is specifically required for preinvasive immunity via recognition of the conserved *P. syringae* effector AvrE1. *CAR1* is highly expressed in guard cells compared to mesophyll cells, and its activation by AvrE1 prolongs stomatal closure initiated by PTI. We also show that the extension of stomatal closure upon activation of CAR1 is dependent on prior PTI-induced stomatal immunity. Finally, we show that CAR1-mediated preinvasive immunity plays a significant role in restricting accessibility of the apoplast to natural isolates of *P. syringae*.

## Results

### AvrE1 induces CAR1-mediated preinvasive immunity

AvrE1 was previously found to trigger CAR1-mediated immunity upon surface inoculation of *A. thaliana* with *P. syringae* ^27^. Here we show that the effectiveness of CAR1-mediated ETI depends on the physical location of the pathogen. First, *A. thaliana* wildtype (WT; natively expressing *CAR1*) was surface inoculated with the tomato and *A. thaliana* pathogen *Pseudomonas syringae* pv. *tomato* PtoDC3000 (hereafter PtoDC3000) carrying the type III secreted effector *avrE1* encoded under its native promoter on the broad host range expression vector pBBR1-MCS-2. As previously shown, PtoDC3000::*avrE1* grew ∼2.5 log10 less than the virulent empty vector control PtoDC3000:::EV at 3 dpi when surface inoculated (Figure 1A). This growth reduction was CAR1-mediated since PtoDC3000::*avrE1* grew ∼2 logs more on the *car1-1* knock-out line than it did on WT plants.

**Figure 1.**
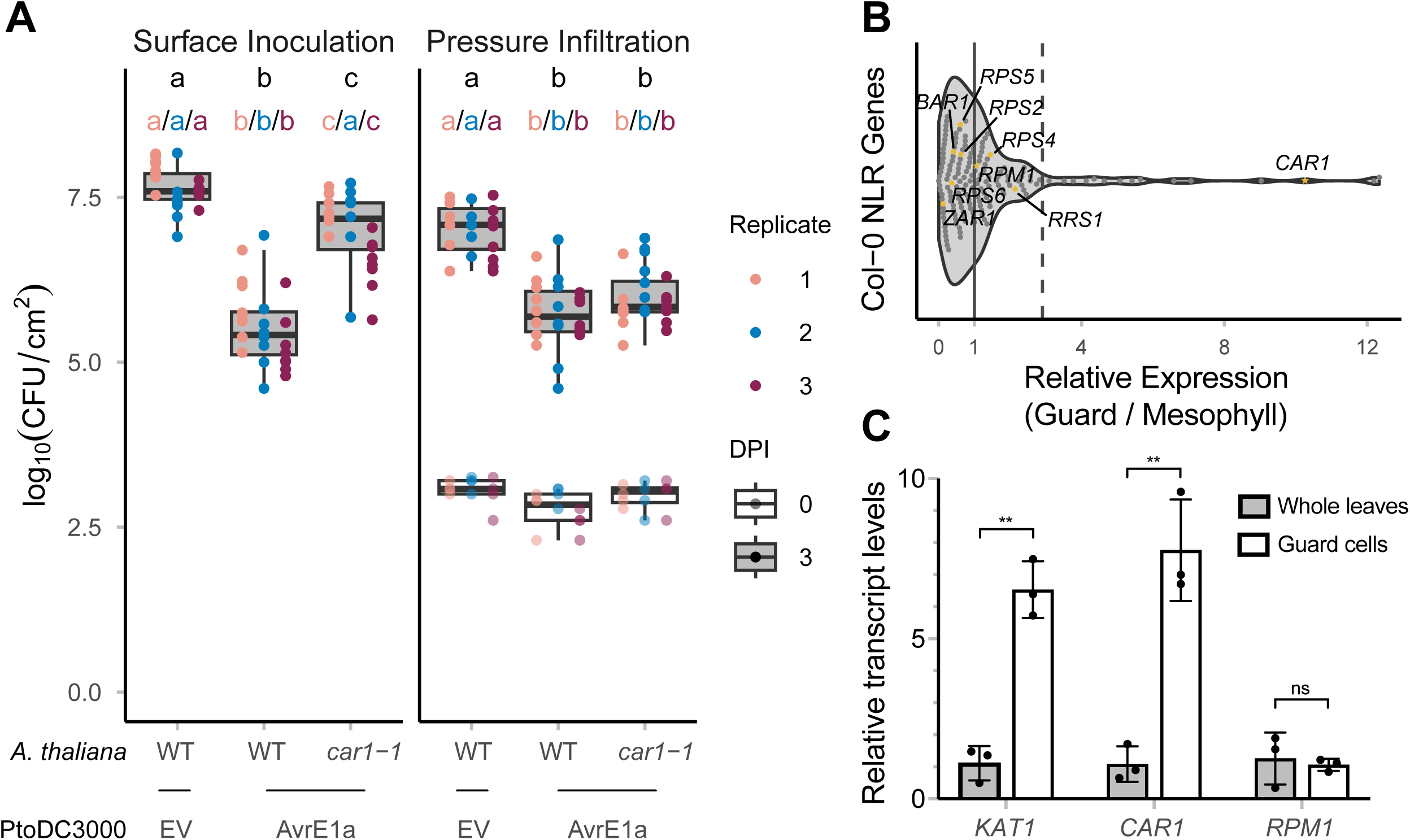
AvrE1 induces CAR1-mediated preinvasive immunity. (A) Comparative *in planta* bacterial growth assays of the virulent PtoDC3000::empty vector (EV) control and PtoDC3000::*avrE1* on *A. thaliana* WT and mutant *car1-1* plants. Infections were done on the same day by diluting the same inoculum for both surface inoculation and pressure infiltration. Each point is a single technical replicate. Different colors represent biological replicates done on different days (n between 6-8 per treatment) with different colored letters indicating statistically significant differences (adjusted p < 0.05, one-way ANOVA followed by post-hoc Tukey-HSD test). Box plots are from pooled biological replicates with pooled statistical differences indicated by black letters. (B) A violin plot displaying the relative gene expression data from the Bio-Analytic Resource ^30^ of 167 NLR or NLR-like genes from the WT reference genome of *A. thaliana*. Each point is the ratio of gene expression in the guard cell tissue versus the mesophyll tissue for each gene. The solid line at 1.0 represents equal expression in both tissue types and the dashed line is the threshold for statistical outliers (1.5 × inter quantile range above the third quartile). Highlighted in orange are the nine identified NLRs that recognize *P. syringae* effectors. Outliers defined as more than 1.5 × inter-quartile range above the third quartile are indicated and listed in Supplementary Table 1. (C) qRT-PCR showing the relative expression of *CAR1* transcript levels in guard cells versus whole leaf extracts. The guard cell marker gene *KAT1* and the NLR *RPM1* (predicted to be equally expressed in both tissue types) were used as positive and negative controls, respectively. Error bars represent standard deviation of three biological replicates with three technical replicates each. ** indicates P < 0.01 and ns indicates P > 0.05 as determined by Student’s t-test.

Remarkably, CAR1-mediated ETI was not observed when the bacteria were injected into the leaf apoplast via pressure infiltration. Here, we found that PtoDC3000::*avrE1* grew ∼1.5 log less than PtoDC3000::EV on WT plants; however, it did not grow to a higher level on *car1-1* plants as observed following surface inoculation (Figure 1A). Overall, AvrE1-triggered immunity was observed in *A. thaliana* by both surface and pressure inoculation, but CAR1 only contributed to immunity following surface inoculation. Because bacteria need to pass through stomata to enter the leaf apoplast after surface inoculation, and this barrier is bypassed by pressure inoculation, we hypothesized that the CAR1 NLR contributes to stomatal immunity.

Further support for this hypothesis came from comparing the relative expression of 167 NLRs or NLR-like genes of *A. thaliana* in guard cells versus mesophyll cells using the Bio-Analytical Resource ^29,30^ (Figure 1B). We found that 15 NLRs (Supplemental Table 1) were significantly overexpressed in guard cells, including *CAR1*, which ranked third with ∼10-fold higher relative expression. To support these observations, we compared the relative expression of *CAR1* in guard cell enriched epidermal extracts versus whole leaf extracts by qRT-PCR (Figure 1C). *CAR1* transcript abundance is significantly higher in the guard cell tissue compared to the whole leaf, similar to the guard cell marker *KAT1* ^31,32^. In contrast, *RPM1* transcript abundance was similar in both extracts as predicted from the Bio-Analytical Resource data (Figure 1C). The enrichment of *CAR1* expression in guard cells supports its proposed stomata-specific role in immunity.

### CAR1-mediated immunity promotes stomatal closure

The PTI response induces stomatal closure approximately one hour following *P. syringae* infection. The phytotoxin coronatine, produced by some strains of *P. syringae*, including PtoDC3000, causes stomata to reopen by approximately four hours post-infection ^16,24^. To determine if CAR1-mediated immunity also plays a role in stomatal closure, we measured stomatal apertures at 0, 1, and 4 hrs after surface inoculation with PtoDC3000::*avrE1* or PtoDC3000::EV (Figures 2A and 2B). Following spray infection with PtoDC3000::EV, we observed an open-close-open stomatal pattern at 0, 1, and 4 hrs, as previously published ^24^. Like PtoDC3000::EV, PtoDC3000::*avrE1* caused stomatal closure at 1 hr after spray infection on WT plants. However, unlike PtoDC3000::EV, the stomata reopening was not observed 4 hrs after infection with PtoDC3000::*avrE1* (Figures 2A and 2B).

**Figure 2.**
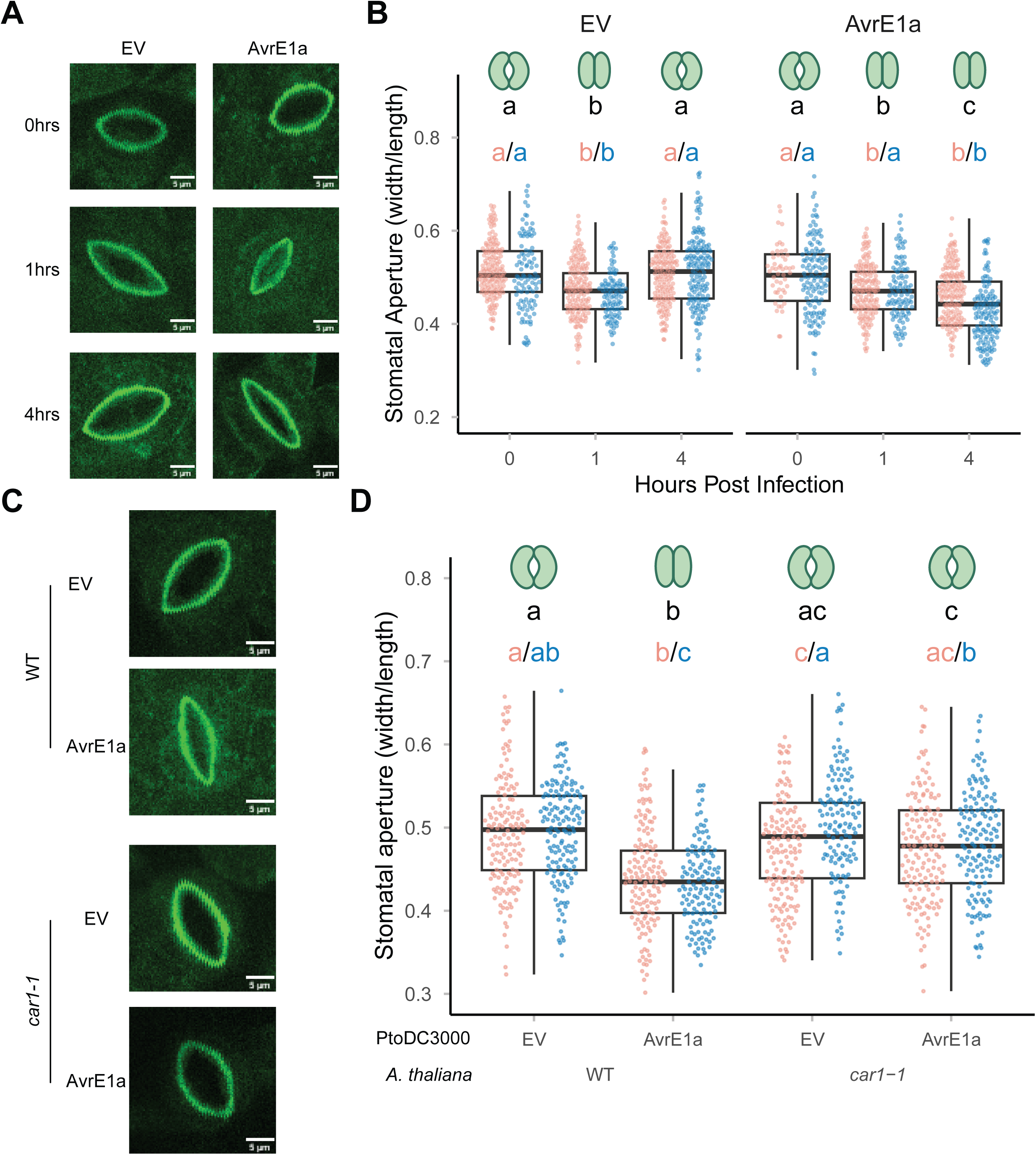
AvrE1 triggers CAR1-mediated stomatal closure. (A) Representative confocal microscopy images of abaxial sides of *A. thaliana* WT plants surface inoculated with PtoDC3000::EV and PtoDC3000::*avrE1*. Tissue was fixed 0, 1, and, 4 hours post-inoculation. The GFP channel was used to visualize rhodamine 6G fluorescence. Scale bars represent 5μm. (B) Stomatal aperture measurements taken from the confocal microscopy images of the GFP channel presented in A (n>100) at 0, 1, and 4 hours post-inoculation. The drawings above each boxplot indicate the expected state of the stomata for each treatment (i.e., open or closed). Each point is an individual stomatal measurement. Different colors represent biological replicates done on different days with different colored letters indicating statistically significant differences (adjusted p < 0.05, one-way Kruskal-Wallis followed by post-hoc Dunn’s test). Box plots are from pooled biological replicates with pooled statistical differences indicated by black letters. (C) Representative confocal microscopy images of abaxial sides of WT and *car1-1 A. thaliana* plants surface inoculated with DC3000::EV and DC3000::*avrE1*. The tissue was fixed 4 hours post-inoculation rhodamine 6G fluorescence was visualized as in A. Scale bars represent 5μm. (D) Confocal microscopy images of the GFP channel presented in C were used to measure stomatal apertures (n>100 per treatment) at 4 hours post-inoculation. The drawings above each boxplot indicate the expected state of the stomata for each treatment (i.e., open or closed). Statistical tests conducted as in B.

To confirm that CAR1 was required to restrict stomatal reopening in response to AvrE1 at 4 hrs after infection, we measured stomatal apertures of WT and *car1-1* plants infected with PtoDC3000::*avrE1* and PtoDC3000::EV (Figures 2C and 2D). Unlike in WT, where we again observed that stomata remain closed following inoculation with PtoDC3000::*avrE1* relative to PtoDC3000::EV, stomatal apertures did not differ following inoculation of the two strains on *car1-1* plants. Thus, the NLR CAR1 sustains stomatal closure until at least 4 hrs after infection in response to the *P. syringae* effector AvrE1.

### CAR1-mediated stomatal closure is contingent on PTI

Given the extensive crosstalk and cooperativity between PTI and ETI, we hypothesized that CAR1-mediated stomatal immunity potentiates the initial PTI-induced stomatal closure. To test this, we infected plants lacking the PRR coreceptor BAK1, which is essential for FLS2 and EFR-mediated PTI responses. *In planta* bacterial growth assays showed that, like CAR1, BAK1 was required for AvrE1-triggered immunity only when bacteria were surface inoculated and not during pressure infiltration (Figure 3A). To further investigate the role of PTI in AvrE1-triggered immunity, we also tested plants lacking the FLS2 gene encoding the PRR that is predominantly required for stomatal immunity against PtoDC3000 ^33^. Similar to BAK1, we observed that FLS2 was required for AvrE1-triggered immunity when bacteria were surface inoculated further supporting the role of PTI in CAR1-mediated immunity (Supplemental Figure 1).

**Figure 3.**
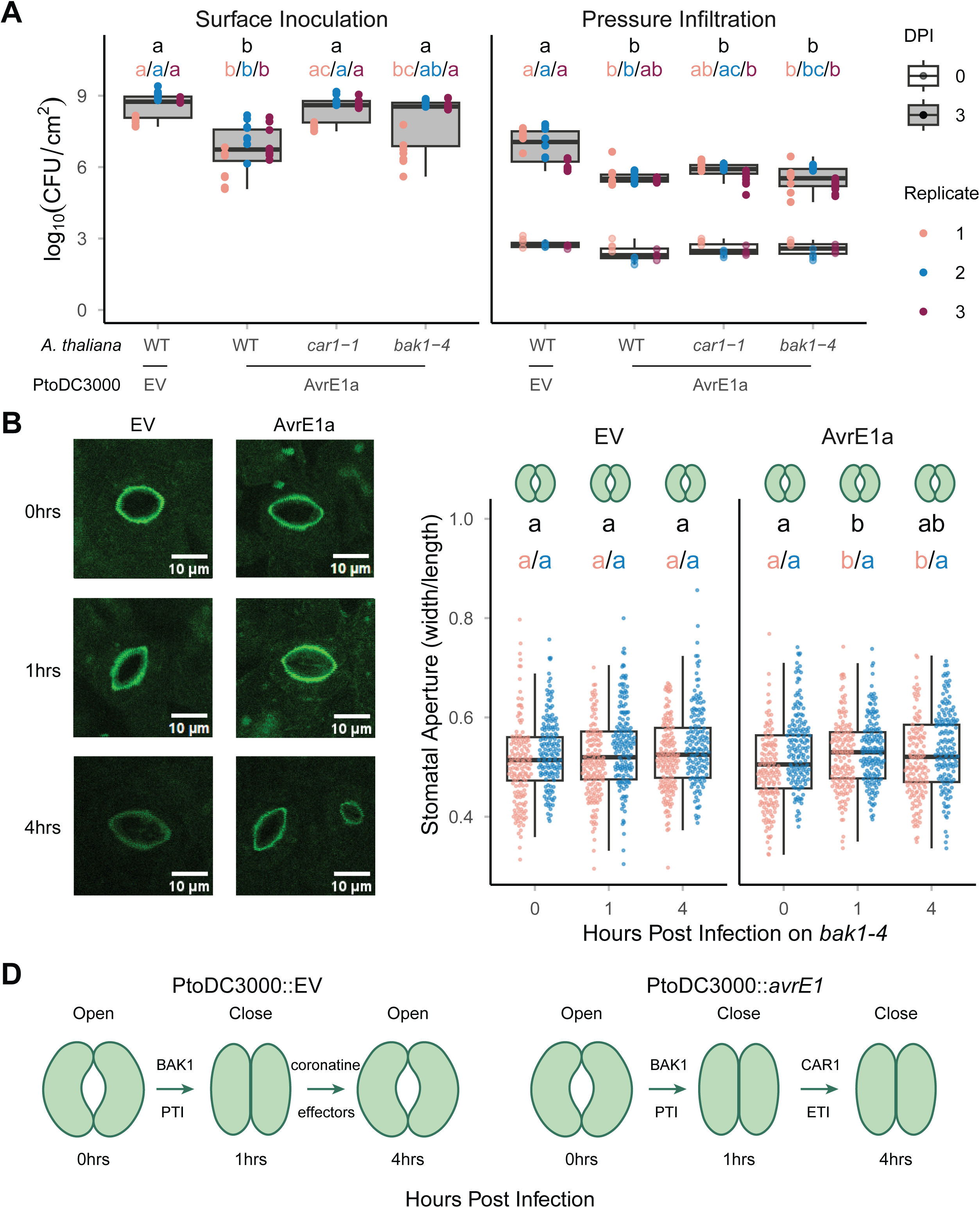
AvrE1-triggered stomatal closure requires the PRR co-receptor BAK1. (A) Comparative in planta bacterial growth assays of the virulent PtoDC3000::EV control and PtoDC3000::*avrE1* on wildtype *A. thaliana* WT, *car1-1*, and *bak1-4* plants. Infections were done on the same day by diluting the same inoculum for both surface inoculation and pressure infiltration. Each point is a single technical replicate, and the different colors represent different biological replicates across three different days (n between 6-8 per treatment) with colored letters representing statistically significant differences (adjusted p < 0.05, one-way Kruskal-Wallis followed by post-hoc Dunn’s test). Box plots are from pooled biological replicates with pooled statistical differences indicated by black letters. (B) Representative confocal microscopy images of abaxial sides of *A. thaliana bak1-4* plants surface inoculated with PtoDC3000::EV and PtoDC3000::*avrE1*. Tissue was fixed 0, 1, and 4 hours post-inoculation. The GFP channel was used to visualize rhodamine 6G fluorescence. Scale bars represent 5μm. (C) Stomatal aperture measurements taken from the confocal microscopy images of the GFP channel presented in B were used to measure stomatal apertures (n>100 per treatment) at 0, 1,and 4 hours post-inoculation. The drawings above each boxplot indicate the expected state of the stomata for each treatment. Each point is an individual stomatal measurement. Different colors represent biological replicates done on different days with different colored letters indicate statistically significant differences (adjusted p < 0.05, one-way Kruskal-Wallis followed by post-hoc Dunn’s test). Box plots are from pooled biological replicates with pooled statistical differences indicated by black letters. (D) Model for stomatal aperture dynamics in response to the effector AvrE1 in the early time points of infection. The series on the left represents stomatal dynamics in response to infection with PtoDC3000; PTI-induced, BAK1-dependent stomatal closure at 1-hour post-infection (hpi) is followed by coronatine/effector-mediated reopening at 4 hpi. The series on the right represents stomatal dynamics following PtoDC3000::*avrE1* infections; PTI-induced, BAK1-dependent stomatal closure at 1 hpi is followed by a CAR1-dependent maintenance of stomatal closure at 4 hpi.

Consistent with the lack of PTI in *bak1-4* plants, stomatal closure was compromised at 1 hr post-infection with PtoDC3000::EV (Figure 3B and 3C). Notably, stomatal closure was compromised at both 1 and 4 hrs after infection of *bak1-4* plants with PtoDC3000::*avrE1* (Figures 3B, 3C). Thus, the CAR1-mediated stomatal closure at 4 hrs post-infection in WT (Figure 2A and 2B) requires prior PTI-induced closure that is dependent on BAK1 and FLS2 (Figure 3D).

### CAR1-mediated immunity contributes to host accessibility of *P. syringae* strains

We previously found that most *P. syringae* strains harbor an ETI-eliciting allele of AvrE1 ^27,34,35^. To test whether CAR1 provides resistance against naturally occurring AvrE1-containing strains, we performed infection assays of wildtype and *car1-1 A. thaliana* with PtoDC3000 and two related strains from phylogroup 1: PtoICMP2844, and Pma4981. Each of these strains carries an ETI-eliciting allele of AvrE1, the coronatine biosynthesis pathway, and no other effectors known to elicit ETI (Figure 4A). Surface inoculation revealed that each strain produced more chlorotic disease symptoms on *car1-1* plants than on WT (Figure 4B and 4C). To quantify pre- and post-invasion infectivity, we compared the *in planta* growth of the three strains following surface inoculation and pressure infiltration on both WT and *car1-1* plants. All three strains showed increased growth on *car1-1* plants relative to WT when surface inoculated, varying from ∼0.5 log CFU/cm^2^ for PtoDC3000 to ∼1 log CFU/cm^2^ for PtoICMP2844 (Figure 4D). In contrast, growth did not differ between WT and *car1-1* for any of the strains following pressure infiltration. These results indicate a generalized role of CAR1 in stomatal immunity against *P. syringae* strains carrying AvrE1.

**Figure 4.**
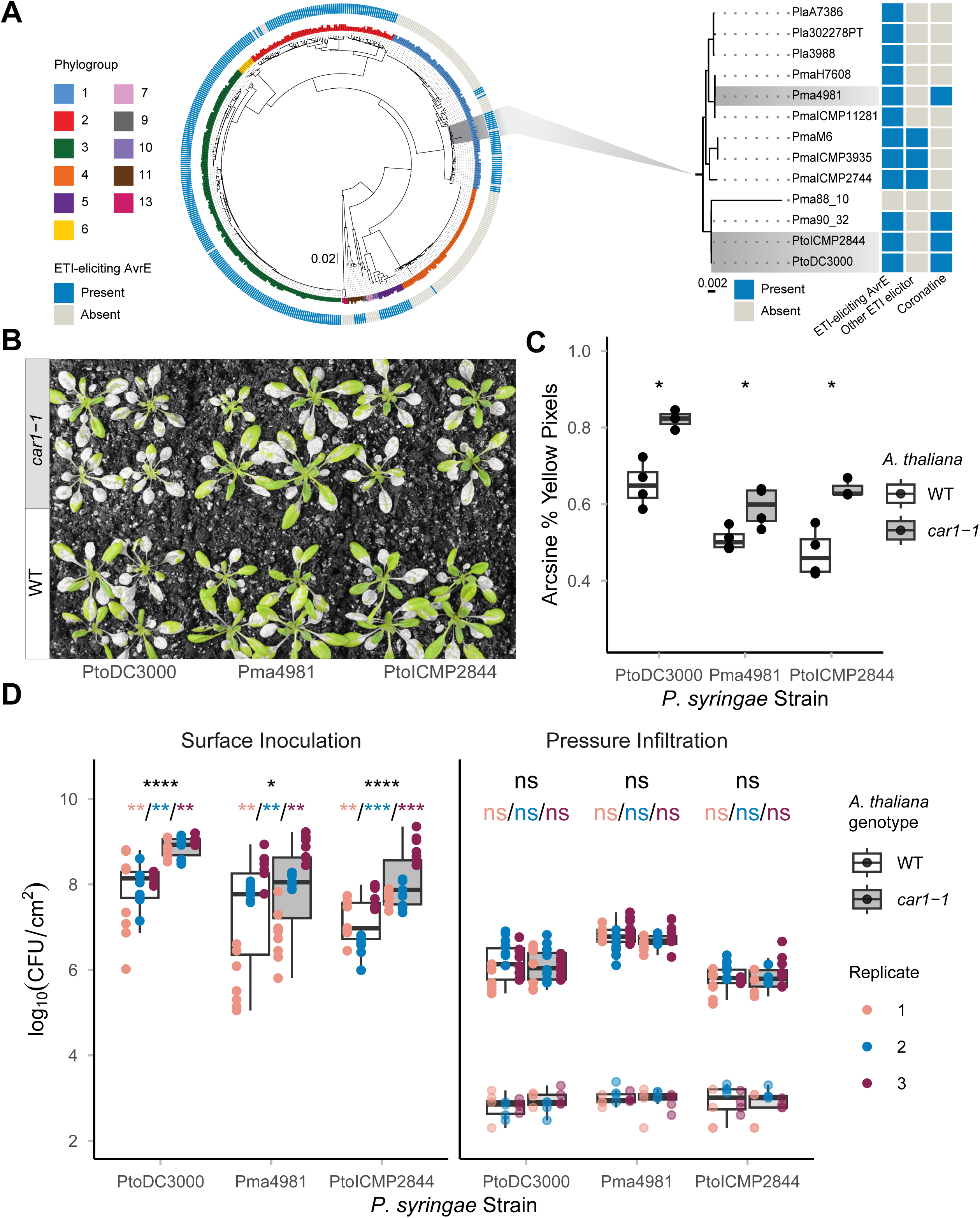
CAR1-mediated immunity contributes to host accessibility of *P. syringae* strains. (A) Left panel: The presence or absence of an ETI-eliciting allele of AvrE1 on *A. thaliana* was mapped onto a core genome maximum likelihood phylogeny of *P. syringae* strains adapted from ^45^. The outer metadata ring is blue for ETI-eliciting AvrE1 presence and grey for absence. Branch tips are coloured by *P. syringae* strain phylogroup, and the scale bar represents genetic distance. Right panel: The clade containing PtoDC3000 was extracted from the core genome tree. The presence or absence of an ETI-eliciting allele of AvrE1 or any other predicted ETI-eliciting effector family ^27^, as well as coronatine (COR), were mapped onto the subtree (data obtained from ^34,45^). PtoDC3000, PtoICMP2844, and Pma4981 (highlighted grey) were selected for investigation since all three possess an ETI-eliciting allele of AvrE1 and COR but no other predicted ETI-eliciting effector. (B) Visual symptoms 7 days post-infection of *A. thaliana* WT and *car1-1* plants surface inoculated with PtoDC3000, Pma4981, or PtoICMP2844. A green-pass filter was applied to enhance the visible differences between healthy and diseased/chlorotic tissue. Four plants were sprayed per treatment per genotype. C) Quantification of plant disease symptoms presented in B (n between 3-4) using PIDIQ; normalized % Yellow Pixels for each plant per treatment ^46^. *P* values: * P < 0.05, ** *P* < 0.01, *** *P* < 0.001, **** *P* < 0.0001, NS *P* > 0.05 (Wilcoxon-Mann-Whitney test). D) Comparative i*n planta* bacterial growth assays of PtoDC3000, Pma4981, and PtoICMP2844 on *A. thaliana* wildtype WT and *car1-1* plants. Infections were done on the same day by diluting the same inoculum for both surface inoculation and pressure infiltration. Each point is a single technical replicate, and the different colors represent different biological replicates (n between 6-8 per treatment). Box plots are from pooled biological replicates with pooled statistical differences indicated by black letters and separate replicate differences indicated by colored letters. Statistical tests and *P* values as described in panel C.

## Discussion

Plants and their foliar pathogens actively fight for control of the stomatal gates. Open stomata enable plants to perform gas exchange and transpiration while providing pathogens access to the leaf interior. In contrast, closed stomata allow plants to block the entry of pathogens, but is also essential for creating an aqueous environment conducive to post-invasive pathogen growth. This study shows that the recognition of the *P. syringae* effector AvrE1 by the *A. thaliana* NLR CAR1 plays a critical role in this battle.

We show that activation of CAR1 by the *P. syringae* effector AvrE1 elicits pre-invasive, tissue-specific ETI that restricts pathogen entry into the leaf tissue by sustaining stomatal closure. This hypothesis is supported by our observations that: (1) CAR1 is required for AvrE1-induced immunity following surface infection but not pressure infiltration, which bypasses stomatal entry, (2) activation of CAR1 sustains stomatal closure, overcoming the reopening driven by coronatine, (3) unlike most NLRs, CAR1 expression is significantly higher in guard cells than mesophyll cells, and finally (4) CAR1 specifically enhances pre-invasive resistance to natural *P. syringae* isolates that possess AvrE1.

Similar to other NLR-mediated immune responses ^36,37^, CAR1-mediated stomatal immunity requires PTI mediated by the PRR-coreceptor BAK1 as well as the PRR FLS2 (Figure 3, Supplemental Figure 1). These findings align with recent demonstrations of cooperativity between ETI and PTI, for example, the necessity of PTI for full ETI against *P. syringae* upon activation of the NLRs RPS2, RPS4, and RPS5 ^4^. The tissue-specific nature of CAR1-mediated immunity demonstrates that this crosstalk can be regulated by cell-type specific expression of an NLR. It remains to be determined whether the potentiation is mutual, with CAR1-mediated immunity reinforcing PTI-mediated stomatal immunity, perhaps by increasing expression of PTI signaling components ^36,37^. It will be interesting to determine whether CAR1 immunity requires other regulatory components known to regulate stomatal apertures such as ABA and the kinase OST1 ^38^.

The AvrE1 – CAR1 interaction beautifully illustrates the “double-edged sword” nature of effectors. On one hand, pre-invasive recognition of AvrE1 by CAR1 enhances PTI-triggered stomatal closure, thereby restricting pathogen entry ^16,39^. On the other hand, following apoplast invasion, AvrE1 is an essential virulence factor ^40^. It promotes a water- and nutrient-rich apoplast by forming water- and solute-permeable channels in the plasma membrane of plant cells and also by promoting abscisic acid synthesis and transport to promote stomatal closure ^24,25,40,41^. A large domain of AvrE1 located on top of the membrane pore within the plant cytosol interacts promiscuously with numerous kinases and phosphatases ^25,42^. Given the predicted cytosolic localization of CAR1, we speculate that it responds to phosphorylation-based signaling initiated by AvrE1.

An intriguing feature of the CAR1-AvrE1 interaction is that CAR1-mediated resistance is not always fully effective. For example, PtoDC3000 is natively virulent on *A. thaliana* despite possessing an ETI-triggering AvrE1 allele. We hypothesize that PtoDC3000 possesses additional effectors that suppress CAR1-mediated immunity. AvrPtoB and HopI are strong candidate suppressors since they have been shown to suppress cell death phenotypes triggered by AvrE1 in *Nicotiana benthamiana* ^43^. Nevertheless, while pathogenic on wildtype *A. thaliana*, PtoDC3000 still exhibits a higher level of growth following surface inoculation on a *car1* mutant line (Figure 4D), indicating that PtoDC3000 is forced to perform a tight balancing act of maintaining AvrE1 virulence function while negating its ETI potential. We can observe the tipping of this balance when we express AvrE1 on a multicopy plasmid and amplify the CAR1-mediated ETI response ^27^. We hypothesize that by increasing AvrE1 dosage, we overwhelm the mitigation strategies that PtoDC3000 possesses to suppress CAR1-mediated recognition. Notably, in addition to CAR1-mediated stomatal immunity, AvrE1 also triggers CAR1-independent, post-invasive immunity (Figure 1A). The nature of post-invasive immunity triggered by AvrE1, perhaps dependent on another NLR, is unknown.

This study identifies an important role for ETI in stomatal immunity. The plant PTI response that rapidly closes stomata to prevent *P. syringae* invasion can be overcome by coronatine and effectors, but activation of CAR1 sustains stomatal closure, thereby temporally extending the effectiveness of this barrier against pathogen entry. It will be interesting to see if stomatal immunity can be driven by stomatal enriched NLRs other than CAR1 and if the conserved NLR CAR1 ^44^ displays a similar tissue-specific response beyond *A. thaliana*.

## Resource availability

All data that support the findings of our study are available in the manuscript or supplementary materials. Code to replicate the analysis in this study is available upon request. Original microscopy images are available upon request.

## Acknowledgments

We thank all the members of the Desveaux and Guttman labs for invaluable discussion and input on this project; B. Laflamme for initial identification of the BAK1 phenotype and J. Cheng for advice on PTI assays. Y. Wang and S. Mukhtar of the Mukhtar lab for help discussion and analysis of sc-RNA seq data. We thank H. Khan of the Provart lab for help with guard cell enrichment. We also thank M. Loranger of the Yoshioka lab for help with qPCR. We also thank members of the Yoshioka and McFarlane labs and the CSB imaging facilities for help with confocal microscopy: R. Goh, M. Loranger, E. Ramirez Rodriguez, and Kenana El Kakouni. D.D. is supported by National Science & Engineering Research Council of Canada Discovery Grants to D.D and D.S.G.

## Author contributions

T.V.A, D.D., and D.S.G designed the project; T.V.A performed all wet lab experiments, T.V.A analyzed and plotted all primary data; Y.L. performed RNA extraction and qRT-PCR analysis; D.Y.W. generated PtoDC3000:*ΔavrE1* mutant; R.A.S. plotted phylogenetic analysis; R.G. and D.M. observed initial pre-invasive phenotype; T.V.A, D.D., and D.S.G wrote the paper; and all authors reviewed and agreed on the manuscript.

## Supplemental information titles and legends

**Supplemental Figure 1.**
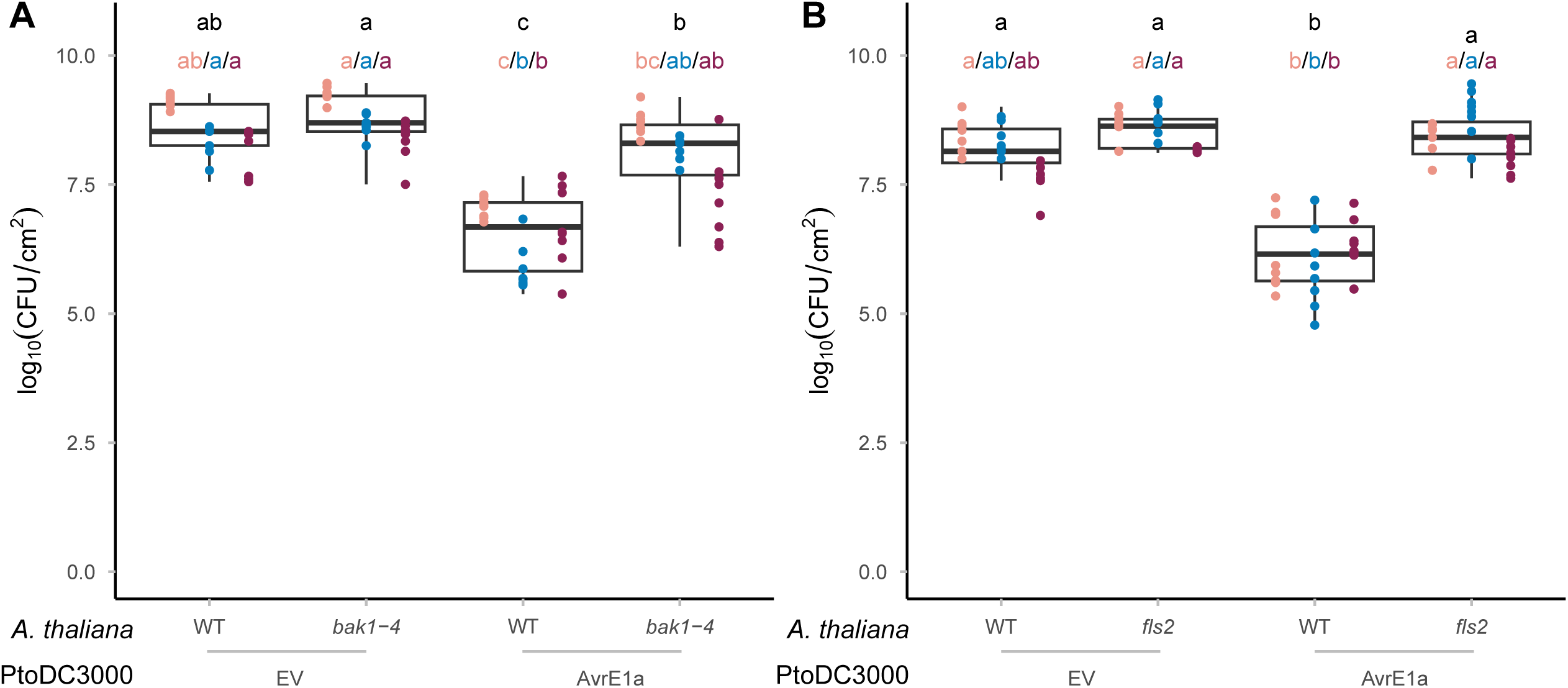
The PRR FLS2 is required for AvrE1-triggered immunity (related to. Figure 3). (A) Comparative in planta bacterial growth assays of PtoDC3000::EV and PtoDC3000::avrE1 surface inoculated on *A. thaliana* wildtype and *bak1-4* WT. Each point is a single technical replicate, and the different colors represent different biological replicates across three different days (n between 6-8 per replicate in treatment) with different colored letters indicating statistically significant differences (adjusted p < 0.05, one-way Kruskal-Wallis followed by post-hoc Dunn’s test). Box plots are from pooled biological replicates with statistical differences indicated by black letters. (B) Comparative in planta bacterial growth assays of PtoDC3000::EV and PtoDC3000::avrE1 surface inoculated on WT and *fls2 A. thaliana* plants. Each point is a single technical replicate, and the different colors represent different biological replicates across three different days (n between 6-8 per replicate in treatment) with different letters indicating statistically significant differences adjusted p < 0.05, one-way Kruskal-Wallis followed by post-hoc Dunn’s test). Box plots are from pooled biological replicates with pooled statistical differences indicated by black letters.

**Supplemental Table 1.** Expression Ratios *A. thaliana* NLR and NLR-Like Loci.

| **Locus ID** | **Name** | **Expression Ratio** |
| --- | --- | --- |
| At5g05400 | LRR and NB-ARC domains-containing disease resistance protein | 12.3 |
| At5g46520 | ACQOS VICTR Disease resistance protein (TIR-NBS-LRR class) family | 12.2 |
| At1g50180 | CAR1 NB-ARC domain-containing disease resistance protein | 10.3 |
| At2g17050 | disease resistance protein (TIR-NBS-LRR class), putative | 9.0 |
| At5g45060 | Disease resistance protein (TIR-NBS-LRR class) family | 7.1 |
| At4g16900 | Disease resistance protein (TIR-NBS-LRR class) family | 6.6 |
| At5g56220 | TN21 P-loop containing nucleoside triphosphate hydrolases superfamily protein | 5.6 |
| At5g41540 | Disease resistance protein (TIR-NBS-LRR class) family | 5.4 |
| At5g46450 | Disease resistance protein (TIR-NBS-LRR class) family | 5.1 |
| At1g15890 | Disease resistance protein (CC-NBS-LRR class) family | 4.6 |
| At3g25510 | disease resistance protein (TIR-NBS-LRR class), putative | 4.4 |
| At5g45220 | Disease resistance protein (TIR-NBS-LRR class) family | 4.1 |
| At5g51630 | Disease resistance protein (TIR-NBS-LRR class) family | 3.8 |
| At5g45050 | ATWRKY16 WRKY16 Disease resistance protein (TIR-NBS-LRR class) family | 3.5 |
| At1g59218 /<br>At1g58848 | RPP13 Disease resistance proteins (CC-NBS-LRR class) family | 3.1 |

NLR or NLR-like genes from *A. thaliana* with an expression ratio between the guard and mesophyll tissue which is more than $1.5 \times$ inter-quartile range above the third quartile relative to all other NLR or NLR-like genes in the Col-0 *A. thaliana* genome.

**Supplemental Table 2.**
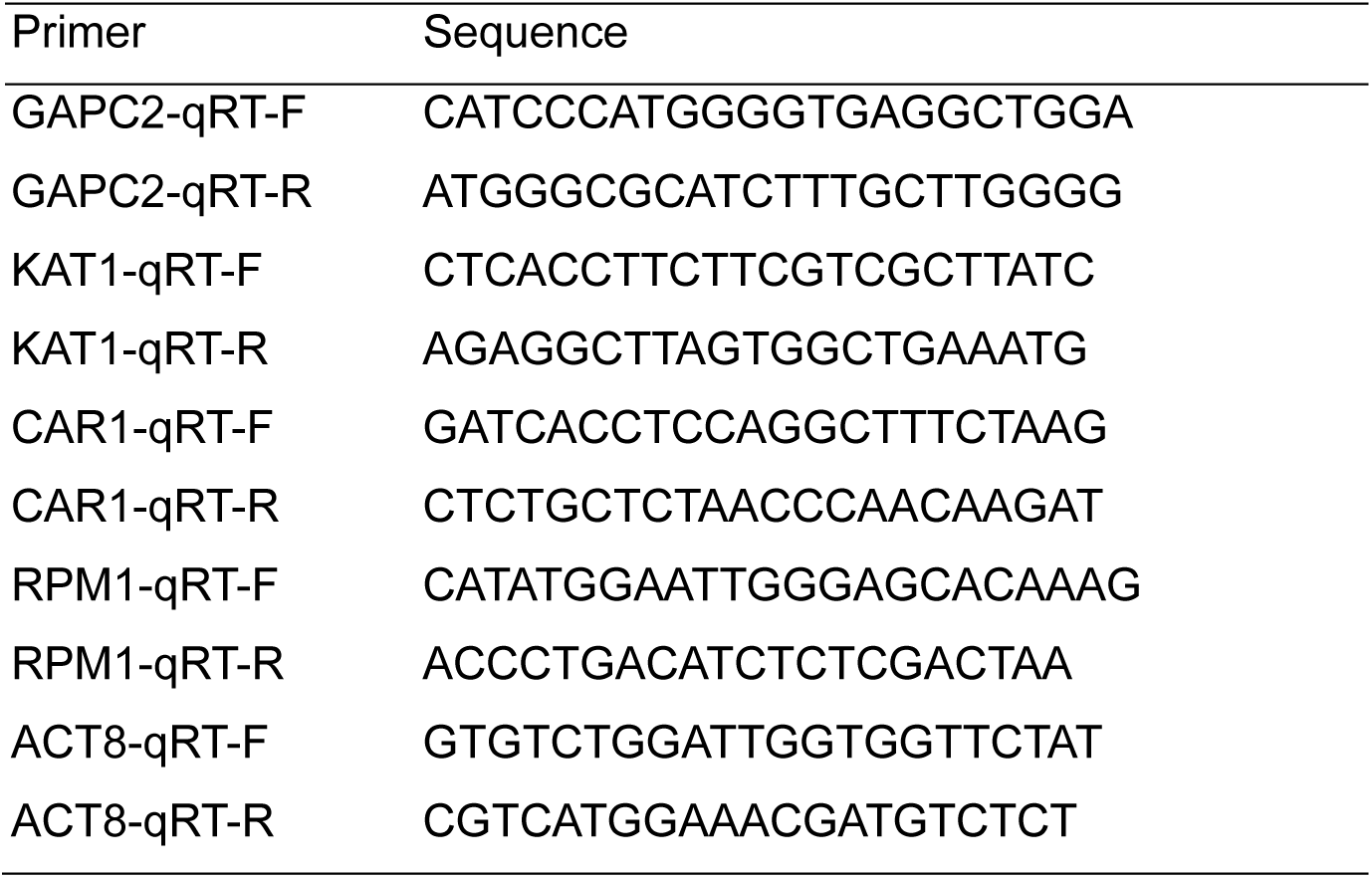
qRT-PCR Primers.

## Methods

### Plant and bacterial growth conditions

All *A. thaliana* plants were grown with a 12-hour photoperiod (130-150 µmol/m^2^s) at 21°C and a 12h dark period at 20°C. Plants were grown in Sunshine Mix 1 soil with regular watering for 24-28 days prior to carrying out surface inoculation or pressure infiltration assays. All *P. syringae* strains were grown at 28°C in King’s B (KB) media. Media was supplemented with the following antibiotics where appropriate: 50 ug/mL of rifampicin (for PtoDC3000, Pma4981, and PtoICMP2844) and/or 50 μg/mL kanamycin (for all PtoDC3000 strains carrying pBBR1-MCS-2 constructs).

### *In planta* bacterial growth assay

(A) *P. syringae* strains were grown overnight on appropriate antibiotics and re-suspended in 10 mM MgSO_4_. Re-suspended strains were diluted to 4x10^8^ CFU/mL (OD 0.8) and 1x10^5^ CFU/mL (OD 0.0002) for surface inoculation and pressure infiltration, respectively. 0.04% Silwet L-77 was added to the surface inoculation suspension. For surface inoculation, 24–28-day-old *A. thaliana* plants were surface infected with roughly 2mL of bacterial suspension per plant using Preval sprayers. Flats were immediately domed to maintain high humidity (∼95%) until the end of the experiment similar to the protocol of Roussin-Léveillée et al. 2022. The ambient room humidity where experiments were conducted fluctuated between 20 and 55%. For pressure infiltration, 24–28-day-old *A. thaliana* plants were pressure-infiltrated into a full leaf with a needleless syringe. Leaves were dried thoroughly after infiltration with a Kimwipe and flats were left undomed until the end of the experiment at ambient room humidity (20-55%).

For bacterial growth assays, four leaf disks (1 cm^2^) from each plant (one per leaf, 8 plants per treatment) were harvested 3-days post-infection. Leaf disks were homogenized using a bead-beater in 1mL of sterile 10 mM MgSO_4_, serially diluted in 96-well plates, and spot-plated on KB supplemented with rifampicin. Plates were incubated for a minimum of 24 hours; the resulting colony counts were used to calculate the number of CFUs per cm^2^ in the leaf apoplast.

### Disease quantification

Plants were surface inoculated with *P. syringae* strains as described in the previous section. Plants were photographed at 7-days post-infection using a Nikon D5200 DSLR camera fitted with a Nikon 18-140 mm DX VR lens. Individual plants were cropped from each image using ImageJ and the resulting cropped photos were analyzed with plant immunity and disease image-based quantification (PIDIQ) ^46^ to determine the amount of chlorotic or yellow tissue.

### Stomatal aperture assay

The stomatal aperture assay was adopted from Roussin-Leveillee *et al.* 2022. 24-28-day old *A. thaliana* plants were surface inoculated with bacterial suspensions of 4x10^8^ CFU/mL (OD 0.8) supplemented with 0.04% Silwet L-77 in 10 mM MgSO_4_. For consistency only leaves from the primary rosette were sampled with developmental leaves 5-8 being preferentially sampled.

Leaves were cut at the base of the petiole and immediately immersed in stomatal fixation solution (4% formaldehyde with 1μM Rhodamine 6G) for 1 minute to fix and stain stomata. A number 3 core was taken from each leaf and cores were mounted in 50% glycerol abaxial side up for imaging. Stomata were imaged using a Leica SP8 confocal microscope under the following settings: excitation at 488 nm, emission 505–545 nm (green fluorescence). The 40X oil immersion objective lens was used to image, and 16-35 tiles were imaged per sample (totalling roughly 50-100 stomates) and automatically stitched together using the Leica Application Suite (LAS X) software. For each experiment stomata were measured off 2-3 different plants per treatment. 2 separate biological replicate experiments were performed in total on different days using fresh inoculum.

### RNA isolation and qRT-PCR analysis

Leaves from 3- to 4-week-old *A. thalian*a WT plants were used for total RNAs isolation. Guard cell-enriched samples were prepared as previously described in ^31,50^. Briefly, for each biological replicate, full rosette of four different plants were harvested and blended in ice-cold deionized water for 1-2 minutes per cycles. Crushed ice was added during blending to facilitate homogenization. Epidermal fragments were filtered through a 210-μm nylon mesh and immediately frozen in grinded liquid nitrogen.

Total RNA from whole leaf tissue and guard cell-enriched epidermal fragments were extracted using TRIzol reagent (Invitrogen) and treated with ezDNase (Invitrogen). Reverse transcription was performed using SuperScript IV VILO (Invitrogen). Quantitative real-time PCR (qRT-PCR) was carried out with a CFX Connect detection system (Bio-Rad) using PowerTrack SYBR Green Master Mix (Applied Biosystems). Relative gene expression data were calculated by modified 2-ΔΔCt method using both *Actin8* (AT1G49240) and *GAPC2* (AT1G13440) as internal reference genes ^51^. All primers used for qRT-PCR are listed in Supplemental Table 2.

